# Population-genomic inference of the strength and timing of selection against gene flow

**DOI:** 10.1101/072736

**Authors:** Simon Aeschbacher, Jessica P. Selby, John H. Willis, Graham Coop

## Abstract

The interplay of divergent selection and gene flow is key to understanding how populations adapt to local environments and how new species form. Here, we use DNA polymorphism data and genome-wide variation in recombination rate to jointly infer the strength and timing of selection, as well as the baseline level of gene flow under various demographic scenarios. We model how divergent selection leads to a genome-wide negative correlation between recombination rate and genetic differentiation among populations. Our theory shows that the selection density, i.e. the selection coefficient per base pair, is a key parameter underlying this relationship. We then develop a procedure for parameter estimation that accounts for the confounding effect of background selection. Applying this method to two datasets from *Mimulus guttatus*, we infer a strong signal of adaptive divergence in the face of gene flow between populations growing on and off phytotoxic serpentine soils. However, the genome-wide intensity of this selection is not exceptional compared to what *M. guttatus* populations may typically experience when adapting to local conditions. We also find that selection against genome-wide introgression from the selfing sister species *M. nasutus* has acted to maintain a barrier between these two species over at least the last 250 ky. Our study provides a theoretical framework for linking genome-wide patterns of divergence and recombination with the underlying evolutionary mechanisms that drive this differentiation.

---

Estimating the timing and strength of divergent selection is fundamental to understanding the evolution and persistence of organismal diversity [1–3]. Genes underlying local adaptation and speciation act as barriers to gene flow, such that genetic divergence around these loci is higher compared to the rest of the genome. However, a framework that explicitly links observable patterns of DNA polymorphism with the underlying evolutionary mechanisms and allows for robust parameter inference has so far been missing [4].

One way of studying adaptive genomic divergence in the face of gene flow is to apply methods for demographic inference to scenarios of speciation [e.g. 5, 6]. This approach allows dating population splits and inferring the presence or absence of gene flow, yet generally does not explicitly account for natural selection [but see 7]. Another approach is to scan genomes for loci that are statistical outliers of divergence among populations. These scans are used to identify candidate loci underlying speciation or local adaptation [e.g. 8, 9], and include the search for so-called genomic islands of divergence [e.g. 10], i.e. extended genomic regions of elevated divergence. Methods of this type can be confounded by other modes of selection as well as demography, and will always propose a biased subset of candidate loci [11, 12].

A third approach is to test for a negative correlation between absolute genetic divergence and recombination rate across the genome [e.g. 13–15]. This approach is based on the prediction that divergence will be higher in regions of the genome where genetic linkage between neutral sites and loci under divergent selection is higher on average [16]. Testing for this pattern of a negative correlation is powerful because it aggregates information across the entire genome and because it is specific to divergent selection with gene flow [17]. However, this approach is purely descriptive and problematic if recombination covaries with a confounding factor, e.g. gene density, that in turn affects the intensity of selection [18].

Here, we develop theory describing the pattern used by this third approach, and a way of inferring the underlying parameters. Our approach explicitly models selection against gene flow and its effect on neutral variation, estimates the strength and timing of selection and gene flow, and filters out the confounding effect of background selection.

## Idea of Approach and Population-Genomic Model

Here, we exploit the genome-wide variation in recombination rate and its effect on genetic divergence. Divergent selection reduces effective gene flow at neutral sites, and this effect decreases with the recombinational distance from the loci under selection. We conceptualize this relationship in terms of the effective migration rate and the expected pairwise between-population coalescence time (Fig. 1A). The latter directly connects to the absolute genetic diversity between populations, a quantity that is readily estimated from DNA sequence data. Our model considers two populations of diploids with effective sizes *N*_1_ and *N*_2_ and non-overlapping generations. In population 1, a balance between one-way gene flow from population 2 at rate *m* per generation and local directional selection is maintained for *τ* generations before the present. In this *migration-selection* (MS) phase (Fig. 1A), selection against maladaptive immigrant alleles acts at an arbitrary number of biallelic loci that we call *migration–selection polymorphisms* (MSPs). At each MSP, one allele is favored in population 1 over the other by an average selection coefficient *s*, while the deleterious allele is introduced by gene flow. We assume additive fitness and no dominance.

**Figure 1.**
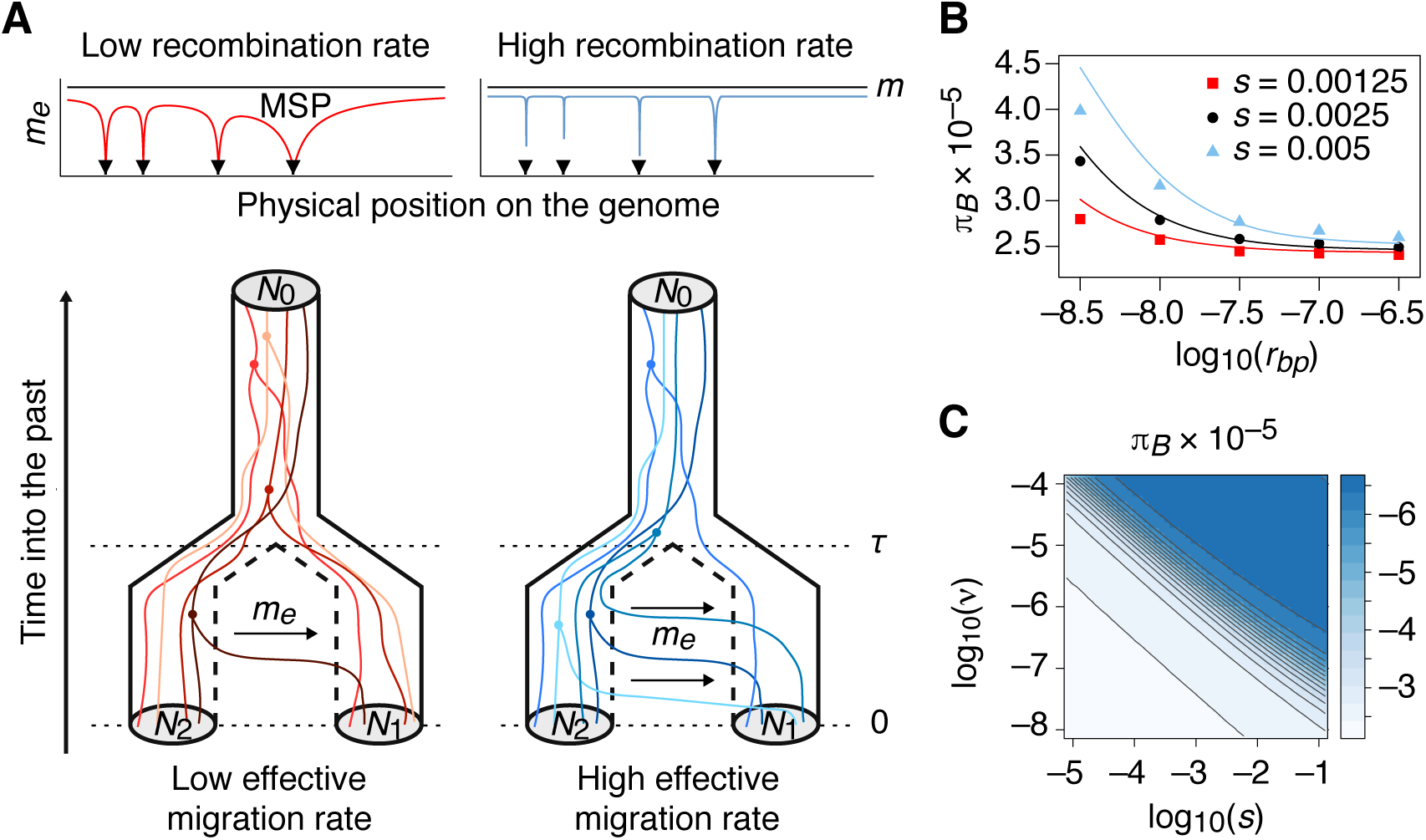
Divergent selection reduces gene flow and increases genetic divergence. *(A)* Selection against locally maladapted alleles at migration–selection polymorphisms (MSPs; black triangles) reduces the effective migration rate *m_e_*. The effect is stronger in regions of low recombination (red, top left) and decreases the probability that lineages sampled in different populations migrate and coalesce. Realizations of the coalescence process are shown in the bottom left for the (MS)P scenario (Fig. S1.1). In regions of high recombination, *m_e_* is higher (blue, top right), such that migration and earlier coalescences are more likely (bottom right). *(B)* The predicted between-population diversity 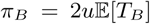 (curves) matches individual-based simulations (dots); error bars (±SE) are too short to be visible. The (MS)M scenario was used with *N*_2_ = 5000, *u* = 10^−9^, *ν* = 2.5 × 10^−7^, *m* = *m*_0_ = 5 × 10^−4^, and *τ* = 4*N*_2_. *(C)* Approximately linear contour lines with slope −1 in the surface of *π*_*B*_ as a function of log_10_(*s*) and log_10_(*ν*) support the compound parameter selection density, *σ* = *sν*. Here, *r*_bp_ = 10^−8^ (1 cM/Mb); other parameters as in *(B)*.

Prior to the MS phase, we assume a *panmictic* (P) phase in an ancestral population of effective size *N*_0_ that starts *τ* generations ago and extends into the past (Fig. 1A). We call this the (MS)P demographic scenario. The P phase can be exchanged for an ancestral *migration* (M) phase with gene flow at rate *m*_0_. Here, we use the (MS)P and (MS)M scenarios to describe our approach. In SI Appendix A we provide extensions to more general scenarios with an intermediate *isolation* phase.

We denote the per-base pair recombination rate by *r*_bp_ and assume that the MSPs occur at a constant rate *ν* per base pair, such that the distance between consecutive MSPs is exponentially distributed with mean 1/*ν* base pairs.

## Average Effective Gene Flow and Selection Density

Selection against maladapted immigrant alleles acts as a barrier to gene flow in the MS phase. At a focal neutral site, the baseline migration rate *m* is reduced to an effective migration rate *m*_e_ [19, 20]. This reduction in effective gene flow increases with the strength of selection at the MSPs, and decreases with their recombinational distance from the focal neutral site (Fig. 1 A; Eq. S1.1 in SI Appendix A). To extrapolate from a given neutral site to the entire genome, we need to average over the possible genomic locations and selection coefficients of the MSPs. For simplicity, we assume an infinite chromosome with a linear relationship between physical and genetic map distance. Given an exponential distribution of selection coefficients, the expected effective migration rate depends on *s*, *ν*, and *r*_bp_ exclusively through *σ*/*r*_bp_, where *σ* = *sν* is the product of the mean selection coefficient times the density of MSPs (SI Appendix A). Note that *σ*/*r*_bp_ has the meaning of a selection density per genetic map unit. For instance, conditioning on two MSPs on each side of an average neutral site, we find 
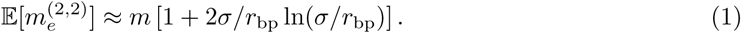

This equation is a good approximation if *σ*/*r*_bp_ ≲ 0.1, i.e. if recombination is at least ten times stronger than selection, at which point effective gene flow is reduced by about 50% (Fig. S1.2 in SI Appendix A). Equation (1) shows that the mean effective gene flow decreases with selection density and increases with recombination rate. Adding increasing numbers of MSPs has a diminishing effect on 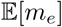 (Fig. S1.2A), so that Eq. (1) captures the essential pattern if *σ*/*r*_bp_ ≲ 0.1. The exclusive dependence of 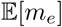 on selection and recombination through the compound parameter *σ*/*r*_bp_ holds for any number of MSPs and so applies to the genome-wide average of *m_e_* (SI Appendix A, Eq. S1.8). Our results imply that doubling the number of MSPs has the same effect on average effective gene flow as doubling the mean selection coefficient. We therefore anticipate that, in practice, *s* and *ν* can be inferred only jointly as *σ* from population-genomic data in our framework.

## Expected Pairwise Coalescence Time With Selection

To facilitate parameter inference from population-genomic data, we phrase our theory in terms of the expected coalescence time of two lineages, one from each population. The expectation of this coalescence time under neutrality, *T_B_*, depends on the baseline migration rate *m* (Table S1.2 in SI Appendix A). We incorporate the effect of selection by substituting the effective migration rate for *m*. Averaging over all possible numbers and genomic locations of MSPs, we obtain 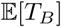, and can predict the between-population diversity *π_B_* as 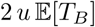, where *u* is the mutation rate per base pair and generation.

To better reflect real genomes, we now assume a finite genome size and define *r_f_* = 0.5 as the recombination rate that corresponds to free recombination, such that MSPs located more than *k_f_* = 1/(2*r*_bp_) base pairs from a neutral site are unlinked. We start by assuming that *ν* is so small that at most a single, nearest-neighboring MSP is linked to any focal neutral site. In the simplest case of the (MS)M scenario with *m*_0_ = *m* (Fig. S1.1C in SI Appendix A), the expected pairwise between-population coalescence time is approximately 
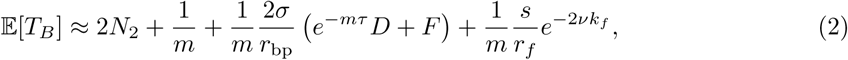

where *D* and *F* depend on *m*, *τ*, and *ν* (Materials and Methods). The first two terms in Eq. (2) are the expectation without selection [21] (Table S1.2). The third and fourth term reflect the increase in coalescence time if the MSP is linked and unlinked to the neutral site, respectively. Importantly, the term accounting for a linked MSP shows that *σ*/*r*_bp_ strongly determines 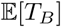, although *s*, *ν*, and *r*_bp_ also enter Eq. (2) independently. Indeed, given *r*_bp_ and in the parameter range where Eq. (2) is a good approximation (i.e. for *ν* ≪ *r*_bp_/*s, m, τ*), the effect of selection on 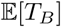 is entirely captured by *σ* (Fig. S1.4 in SI Appendix A). For details and other demographic scenarios, see SI Appendix A. In this simplified model, the effect of all other MSPs is absorbed by *m* as a genome-wide reduction in gene flow that is independent of recombination.

In practice, we want to explicitly account for all MSPs possibly present in the genome, as well as for the average physical chromosome length. Finding 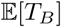 in this more realistic setting amounts to averaging over all possible numbers and genomic locations of the MSPs. We wrote a C++ program to do this integration numerically (SI Appendix A). The result agrees well with individual-based forward simulations (Fig. 1B; Fig. S1.5 in SI Appendix A). As with a single MSP, if *r*_bp_ is given, 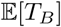 depends on *s* and *ν* effectively only through the selection density *σ* (Fig. 1C). In fact, returning to the idealizing assumption of a global linear relationship between physical and genetic map distance (i.e. *r_f_* → ∞), we show that this dependence holds exactly (SI Appendix A, Eq. S1.68). This finding corroborates *σ* as a key parameter and natural metric to quantify genome-wide divergent selection in the face of gene flow.

## Application to Mimulus guttatus

We developed an inference procedure based on our theory and applied it to two datasets from the predominantly outcrossing yellow monkeyflower (*Mimulus guttatus*), an important model system for speciation and local adaptation [22]. For both datasets, we fit the model with multiple MSPs to the empirical relationship between recombination rate (*r*_bp_, estimated from a linkage map) and putatively neutral between-population diversity (*π_B_*, estimated from 4-fold degenerate coding sites), after correcting the latter for genomic correlates and divergence to the outgroup *M. dentilobus* (SI Appendix B). Our procedure computes the sum of squared deviations (SSD) across genomic windows between these observed values of *π_B_* and those predicted by our model, given the estimate of *r*_bp_ for each window and a set of parameter values. Minimizing the SSD over a large grid of parameter values, we obtained point estimates for the selection density (*σ*), baseline migration rate (*m*), and duration of the MS phase (*τ*). We estimated 95% non-parametric confidence intervals (CIs) for the parameters by doing a block-bootstrap over genomic windows (SI Appendix B). For both datasets, we explored two alternative demographic scenarios, but focus here on the one that provided more plausible parameter estimates and tighter 95% CIs. Unless otherwise stated, we only report results obtained with genomic windows of 500 kb because results for windows of 100 and 1000 kb were very similar (SI Text 2).

## Accounting for background selection

In *M. guttatus*, pericentromeric regions are gene-poor and characterized by low recombination rates, which results in a genome-wide positive correlation between gene density and recombination rate [14] (Fig. S4.4 in SI Text 2). Such a correlation could attenuate or even reverse the positive correlation between diversity and recombination rate otherwise expected under background selection (BGS) and other modes of selection at linked sites, because the strength of such selection is in turn expected to be positively correlated with gene density [18]. This effect could create a false signal of selection against gene flow, as the increased coalescence rate within populations in gene-dense regions could produce a negative correlation between *r*_bp_ and *π_B_*. Comparing tests of a partial correlation between *r*_bp_ and diversity with and without gene density as a covariate, we found that this effect might be present in our first dataset (SI Text 2; File S4.1). Therefore, we first fit a BGS model to the genetic diversity *within* source populations by allowing the effective population size (*N*_2_) to vary as a function of gene density and *r*_bp_. We then incorporated BGS into our migration-selection inference procedure, using these predicted relationships between *N*_2_, gene density, and *r*_bp_ (SI Appendix B). This procedure filters out the effect of BGS since, with unidirectional gene flow, BGS in the *source* but not the *focal* population may affect *π_B_* (Eq. 2).

## Adaptive divergence maintained in the face of gene flow

*Mimulus guttatus* can be found growing on serpentine soils throughout its range [25, p. 4]. While the mechanism and molecular basis of this adaptation are unresolved [26], strong differences in survival on serpentine soil exist between serpentine and non-serpentine ecotypes [25]. To see whether there was a population-genomic signal of local adaptation, we used whole-genome pooled-by-population sequencing of 324 individuals collected from two pairs of geographically close populations growing on and off serpentine soil in California (the serpentine dataset; Fig. 2A; SI Text 1). We inferred the strength of selection in serpentine populations (REM and SLP) against maladaptive immigrant alleles from the geographically proximate off-serpentine population (SOD and TUL, respectively), using the latter in each pair as a proxy for the source of gene flow. These pairs of geographically close serpentine × off-serpentine populations are genetically less diverged than any other population pair (Fig. 1A; Fig. S4.1 in SI Text 2). Since we observed a strong signal of BGS in all populations (SI Text 2), we corrected for this signal when fitting our migration–selection model to the data (SI Appendix B). We found that the conditional surface of the −SSD (holding *m* and *τ* at their point estimates) showed a pronounced ridge for *s* and *ν*, with the 95% confidence hull falling along this ridge (Fig. 2B). With parameters on a log_10_ scale, the slope of this ridge is −1, nicely confirming our theoretical result that *s* and *ν* should be estimated jointly as their product, the selection density *σ*. We therefore adjusted our inference procedure to jointly infer *m*, *τ*, and *σ* instead of *m*, *τ*, *s*, and *ν* (SI Appendix B). This adjustment resulted in profile −SSD surfaces for *σ* and *m* with a unique peak and tight confidence hulls (Fig. 2C).

**Figure 2.**
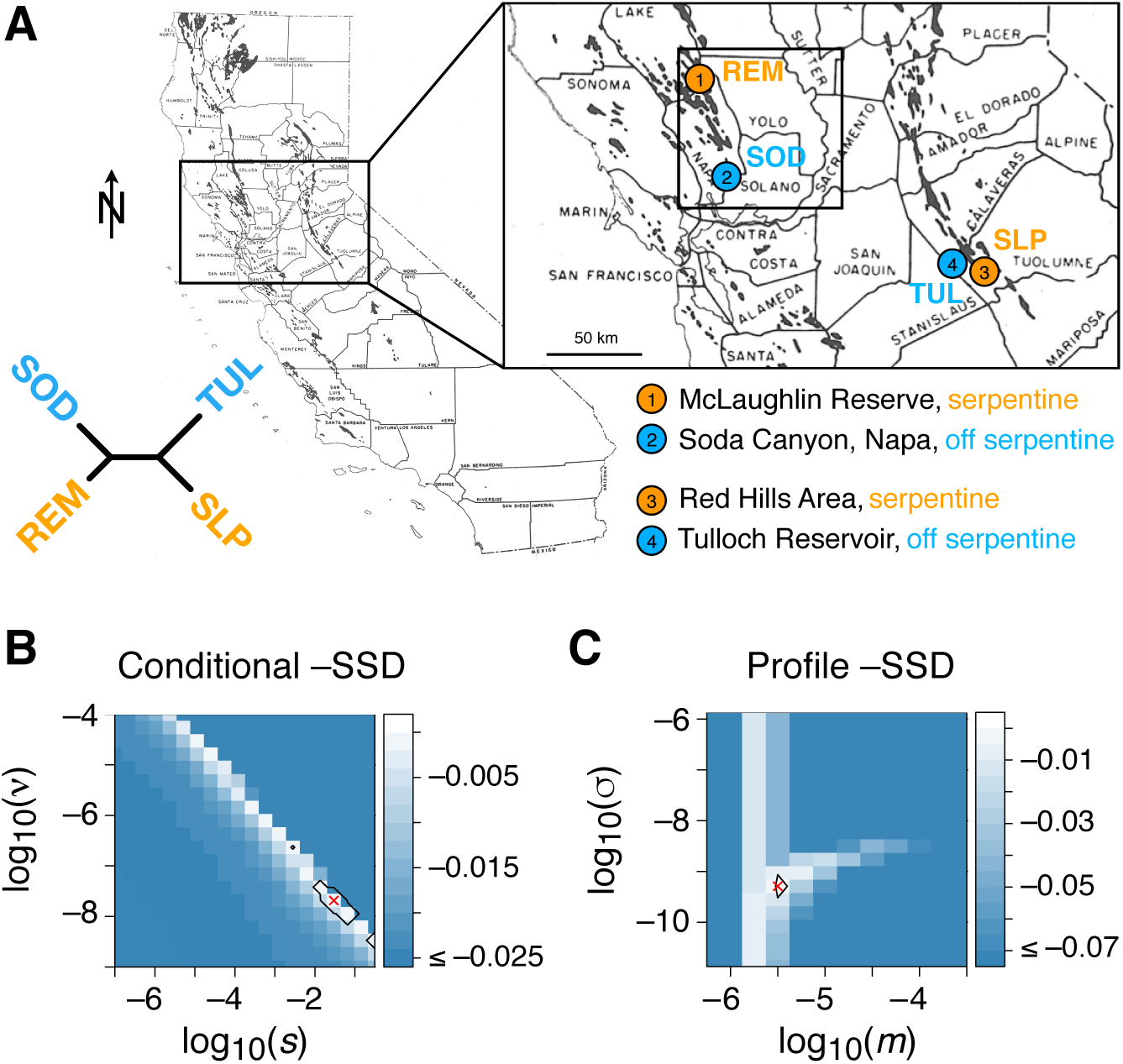
Geographic context of serpentine dataset and quasi-likelihood surfaces. (*A*) Sampling sites in California, USA (modified with permission from [23]), and unrooted population phylogeny based on linearized genetic divergence [24]. *(B)* The negative sum of squared deviations (−SSD) for the selection coefficient s and the genomic density ν of MSPs, conditional on estimates of *m* and *τ*. The ridge with slope −1 confirms the compound parameter selection density, *σ* = *sν*. A cross denotes the point estimate and black hulls the 95% bootstrap confidence area. *(C)* Joint profile surface of the −SSD for the baseline migration rate *m* and the selection density *σ*, maximized over *τ*. Results are shown for the population pair REM × SOD under the (MS)M scenario with genomic windows of 500 kb.

For both serpentine × off-serpentine pairs, we found a strong genome-wide signal of divergent selection against gene flow, with point estimates for *σ* of about 8.3 × 10^−4^ and 4.8 × 10^−4^ per megabase (Mb) in REM × SOD and SLP × TUL, respectively, and tight 95% CIs (Fig. 3A, C; File S4.2). Given an assembled genome size of about 320 Mb for *M. guttatus*, this selection density would, for instance, be consistent with about 300 MSPs, each with a selection coefficient on the order of 10^−4^ to 10^−3^. The impact of this selection on genome-wide levels of polymorphism is reflected in an increase in the effective migration rate (*m_e_*) with higher recombination rate (Fig. 3B, D). The 95% CI of the relative difference (*δ_m_e*) between the maximum and minimum of *m_e_/m* clearly excludes 0 (Fig. 3B, D; SI Appendix B), indicating a partial shutdown of gene flow due to selection. According to our estimates of *m*, selection maintains this divergence against a baseline rate of gene flow of about 6.6 × 10^−6^ in REM × SOD and 3.5 × 10^−6^ in SLP × TUL (Fig. 3B, C). Given the estimated effective population sizes of REM and SLP (SI Text 1), these rates of gene flow imply about 3.8 and 2.1 diploid immigrants per generation, respectively.

**Figure 3.**
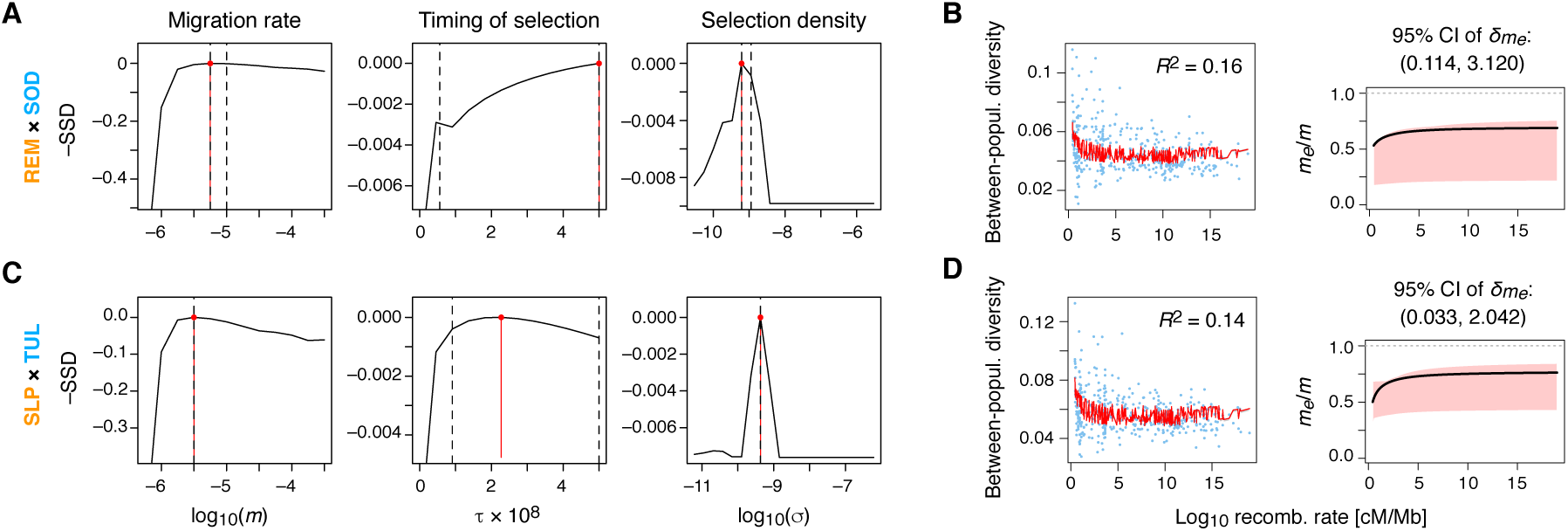
Parameter estimates and model fit for the serpentine dataset. (*A, C*) Profile curves of the quasi-likelihood (−SSD) for each parameter, maximizing over the two remaining parameters, for the serpentine × off-serpentine comparisons REM × SOD (*A*) and SLP × TUL (*C*) (Fig. 2). Vertical red and black dashed lines indicate the point estimate and 95% bootstrap confidence intervals (CIs), respectively. (*B, D*) Raw data (blue dots) and model fit (red curve) with 95% CI (gray shading). The corresponding ratio of the effective to the baseline migration rate is shown on the right (red shading: 95% CI). The 95% CI of the distribution of the relative difference between the maximum and minimum *m_e_* across all bootstrap samples, 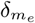, is given on top. Other details as in Fig. 2B-C. For other population pairs, see Fig. S4.17 in SI Text 2.

We had little power to infer precise point estimates for *τ*, but lower bounds of the 95% CIs were around 10 Mya. It seems unlikely that the two ecotypes persisted for so long and so our parameter estimates should be interpreted as a long-term average over a potentially more complex scenario.

To assess if the selection against gene flow we found is specific to serpentine × off-serpentine comparisons (REM × SOD, SLP × TUL), we also fit our model for the two long-distance off-serpentine × off-serpentine configurations (SOD × TUL, TUL × SOD) and the long-distance serpentine × off-serpentine pairs (REM × TUL, SLP × SOD). Interestingly, we inferred selection densities, durations of the MS phase, and migration rates on the same order as those estimated for the short-distance serpentine × off-serpentine comparisons (Fig. S4.17 in SI Text 2; File S4.2). The signal we detect may therefore have little to do with local adaptation to serpentine *per se*, and not be specific to the history of particular pairs of populations. This is corroborated by the fact that, when pooling all non-focal populations to a joint source of gene flow, we observed a similar, if not even stronger, signal of selection against migrants (Fig. S4.18 in SI Text 2; File S4.2). Given the long time *τ* over which this selection appears to have acted, our estimates may reflect adaptive divergence between *M. guttatus* populations in response to locally varying conditions other than serpentine soil [e.g. 27–29]. Our results could also imply that adaptation to serpentine has a simple genetic basis, as our approach only has power to detect a signal that is due to polygenic divergent selection acting across the entire genome.

## Persistence of species barrier to M. nasutus

Where *M. guttatus* has come into secondary contact with *M. nasutus*, a selfing sister species, hybridization occurs despite strong reproductive barriers [30]. A previous genome-wide analysis identified large genomic blocks of recent introgression from *M. nasutus* into sympatric *M. guttatus* populations [14]. Using 100-kb genomic windows, this previous study also found a negative correlation between absolute divergence (*π_B_* = *π*_Gut×Nas_) and recombination rate (*r*_bp_) in sympatric but not allopatric comparisons, as would be expected if there was selection against hybrids. Reanalyzing these data (the GutNas dataset; SI Text 1) we replicate this pattern of correlation. However, if we included gene density as a covariate, all previously negative partial correlations between *r*_bp_ and *π*_B_ became non-significant (Fig. S4.12A in SI Text 2). This observation might indicate that the positive correlation between recombination and gene density could have camouflaged an underlying signal of BGS in the source population, as was the case with the serpentine dataset above. To test this hypothesis, we fit a model of BGS in *M. nasutus*, but found no evidence for BGS (SI Text 2) [cf. 14].

Applying our method to 100-kb windows, we indeed found a significant signal of selection against hybrids for one sympatric pair (CAC × Nas;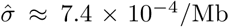), yet no signal in the other (DPR × Nas). We also inferred significant selection against gene flow in one of the allopatric comparisons (SLP × Nas; 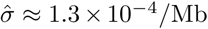). The signal of selection in SLP × Nas could be due to the fact that, although allopatric, SLP is geographically close to M. nasutus populations [14]. We might therefore be detecting selection against ongoing gene flow over a longer distance, or against past gene flow that has stopped only recently. Since levels of recent introgression are much lower in SLP than in the sympatric populations (AHQ and CAC) [14], the second explanation is more plausible. Indeed, repeating our analyses with blocks of recent introgression excluded, we found that the signal of selection against hybrids remained for SLP × Nas, but disappeared for sympatric comparisons (Fig. S4.12B; Fig. S.4.21 in SI Text 2).

Our estimates of *m* for CAC and SLP imply that selection maintains the species barrier against a baseline migration rate of about 10^−6^, i.e. 1.0 and 0.7 diploid introgressing genomes per generation, respectively (Fig. S4.19 in SI Text 2; File S4.3). With blocks of recent introgression excluded, our point estimate of *m* obtained with 100-kb windows dropped by a factor of 2.8 for CAC × Nas (File S4.3), consistent with the removal of a substantial part of recently introgressed DNA. In contrast to the serpentine dataset, our results for the GutNas dataset were sensitive to the choice of window size. For windows of 500 and 1000 kb, the uncertainty in parameter estimates was higher (Fig. S4.20, S4.22 in SI Text 2; File S4.3).

With 100-kb windows and blocks of recent introgression included, lower 95%-CI limits for τ were all above 250 kya. Point estimates were between about 550 kya (AHQ × Nas) and 1.6 Mya (DPR × Nas) (File S4.3). These estimates are somewhat above a previous estimate of 196 kya for the divergence time between *M. guttatus* and *M. nasutus* [14]. Our older estimates of *τ* are compatible with divergent selection acting already in the ancestral, geographically structured, *M. guttatus* clade [14].

## Discussion

The genomes of incompletely isolated species and locally adapted populations have long been thought of as mosaics of regions with high and low divergence [31, 32]. This pattern is due in part to variation in effective gene flow along the genome, created by an interaction of divergent selection and recombination [33, 34]. The recent explosion of genome-wide DNA sequencing data allows us to directly observe this mosaic, and has spurred theoretical and empirical studies aiming to better understand the mechanisms underlying local adaptation and speciation [e.g. 35–38]. Yet, an explicit, model-based framework linking observed genome-wide patterns of divergence with the underlying mechanism has hitherto been missing.

Here, we developed such a framework by merging the concept of effective migration rate with coalescence theory. We showed that a genome-wide negative correlation of between-population diversity with recombination rate [14, 17] can be described by the compound parameter ‘selection density’, such that very different genomic mosaic patterns are compatible with the same aggregate effect of divergent selection and gene flow: a large number of weak genetic barriers to gene flow (MSPs) is equivalent to a much smaller number of strong barriers. Our approach is most sensitive to polygenic selection, and therefore complements existing genome scans for empirical outliers of population divergence [39–42], which tend to identify only strong barriers to gene flow. It also provides a better null model for such genome scans, as outliers could be judged against the appropriate background level of divergence.

Our approach is inspired by earlier work exploiting the genome-wide relationship between recombination rate and genetic diversity within a population for quantitative inference about genetic hitchhiking [43, 44] and background selection (BGS) [45, 46]. In fact, we have used an established model of BGS [47, 48] to filter out any confounding effect of BGS and gene density in a first step, before fitting our model of divergent selection against gene flow to the relationship between recombination rate and diversity between populations.

We have assumed that MSPs occur at a constant rate *ν* along the genome. This assumption could be relaxed by making *ν* depend on the functional annotation of genomes, e.g. exon coordinates, which might allow *ν* and *s* to be estimated separately [see 49]. We explored this solution heuristically for the serpentine dataset by setting *ν* proportional to the local density of exonic sites. Point estimates of *σ* were on the same order as before (10^−10^ to 10^−8.25^). Yet, the 95% CIs became much wider and the variance explained (*R*^2^) dropped from about 15 to 5%. This reduced goodness of fit might suggest that selection against gene flow is not acting exclusively on coding (exonic) variation.

Our model currently does not account for the clustering of locally adaptive mutations arising in tight linkage to previously established MSPs, and the synergistic sheltering effect among MSPs that protects them from being swamped by gene flow [20, 50]. If accounted for, this clustering would lead to an even more pronounced uptick of between-population diversity in regions of low recombination. Therefore, one might be able to use deviations from our current model in regions of low recombination as a way of detecting the presence of clustering in empirical data. At the very least, our parameter estimates would indicate whether and in what genomic regions one should expect clustering of MSPs to have evolved.

An inherent limitation of our approach is that enough time must have passed for between-population divergence to accumulate. Otherwise, there is no power to detect variation in divergence among genomic regions. This limitation constrains the temporal resolution of our method, in particular if the duration of the migration–selection (MS) phase is short, or if strong reproductive isolation evolved so quickly that gene flow was completely and rapidly reduced across the entire genome. Another potential limitation is a relatively low resolution to infer the duration of the MS phase. A genome-wide negative correlation of recombination rate with between-population diversity will persist for a long time even after gene flow has come to a complete halt, because subsequent neutral divergence will just add uniformly to the existing pattern. Our inference approach should therefore still provide good estimates of the strength of selection and gene flow even after speciation has completed, as long as these estimates are interpreted as averages over the inferred time *τ*. In this sense, our approach is likely robust to the specifics of the most recent demographic history of the populations or species of interest. To better resolve the timing of events, we suggest using the additional information contained in the entire distribution of pairwise coalescence times (and pairwise sequence differences), rather than relying on their mean, as we currently do.

The opposing roles of gene flow and selection in speciation and local adaptation have a long and contentious history in evolutionary biology and population-genetics theory [3, 51]. We antic-ipate that the type of genome-wide quantitative inference developed here, applied to the growing amount of whole-genome polymorphism and recombination data, will help to resolve how gene flow is constraining divergent selection.

## Materials and Methods

In Equation (2), *D* = Ei[(1 − *g_f_*)*mτ*] − Ei[(1 − *g*_o_)*mτ*] and *F* = Ei[−*g*_o_*mτ*] − Ei[−*g_f_ mτ*] + Ei[−*νk_f_*] − Ei[−*νk_o_*], where 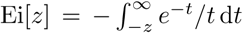 is the exponential integral. Here, *g_f_* = [1 + *s*/*r_f_*]^−1^ and *g_o_* = [1 + *s*/(*k_o_r_bp_*)]^−1^ are the contributions to the gff if the MSP is unlinked (*k*_1_ = *r_f_*/*r_bp_*) or fully linked (*k*_1_ = *k*_o_, 0 < *k*_o_ ≲ 1/[*r*_bp_*τ*]); *k*_o_ is a small positive lower limit for the physical distance to the MSP. For details of our model, theory, and individual-based simulations, see SI Appendix A. Statistical data analyses, bias corrections, and the inference procedure are described in detail in SI Appendix B. The *Mimulus* datasets (sampling design, DNA sequencing, quality filtering), and the linkage map are discussed in SI Text 1. For complementary results, including tests of partial correlation between diversity and recombination rate as well as the inference of background selection, see SI Text 2.

## Acknowledgments

We thank Ben Blackman for help with collections; and Yaniv Brandvain, Lex Flagel, Amanda Kenney, Andrea Sweigart, Kevin Wright, Chenling Xu, and members of the G.C., Ross-Ibarra, and Schmitt laboratories at UC Davis for discussions and comments that improved our study. This work was supported by the National Institute of General Medical Sciences of the National Institutes of Health under grants NIH RO1GM83098 and RO1GM107374 (to G.C.); National Science Foundation (NSF) Grant 1354688 (to J.H.W.); NSF Grant 1353380 (to G.C.); NSF DDIG Grant 1110753 (to J.P.S.); and Swiss National Science Foundation Advanced Postdoc. Mobility Fellowship P300P3_154613 (to S.A.). The computational results presented have been achieved in part using the Vienna Scientific Cluster.

